# The LUX Score: A Metric for Lipidome Homology

**DOI:** 10.1101/013847

**Authors:** Chakravarthy Marella, Andrew E Torda, Dominik Schwudke

## Abstract

**Motivation:** We propose a method for estimating lipidome homologies analogous to the ones used in sequence analysis and phylogenetics.

**Results:** Algorithms were developed to quantify the structural similarity between lipids and to compute chemical space models of sets of lipid structures and lipidomes. When all lipid molecules of the LIPIDMAPS structure database were mapped in such a chemical space, they automatically formed clusters corresponding to conventional chemical families. Homologies between the lipidomes of four yeast strains based on our LUX score reflected the genetic relationship, although the score is based solely on lipid structures.

**Availability:** www.lux.fz-borstel.de

## 1 INTRODUCTION

A lipidome can be an indicator of health, disease, stress or metabolic state. Using model organisms, the role of lipid metabolism has been studied in diseases such as diabetes, metabolic syndrome, neurodegeneration and cancer (Yetukuri *et al.*, 2007; Kühnlein, 2012; Subramanian *et al.*, 2013; Hindle *et al.*, 2011; Lopez and Scott, 2013; Kiebish *et al.*, 2008). To this end, lipidomes from yeast and fruitfly have been characterised (Ejsing *et al.*, 2009; Guan *et al.*, 2010; Carvalho *et al.*, 2012; Klose *et al.*, 2012; Guan *et al.*, 2013) enabling one to identify fundamental lipid metabolic processes (Lam and Shui, 2013; Peng and Frohman, 2012). However, a critical question remains: How relevant are changes in the lipidome of a biological model for understanding human physiology if these lipids are not present in humans?

For example, it would be a challenge to relate differences in lipid metabolism in *D. melanogaster* or *S. cerivisae* to human biochemistry. One would only have to look at their differing sphingolipid compositions (Kraut, 2011). In humans, sphingomyelins (SM) are highly abundant, but they are basically absent in the fruitfly.

Furthermore, drosophila sphingolipids have a shorter sphingoid alkyl chain (C14), but their amide bond fatty acids are usually longer than those in humans.

Homology measures for genes and protein sequences are well established. The theme in this work is the development of similar metrics for lipidome homology. We started by converting lipid structures to Simplified Molecular Input Line Entry Specification (SMILES) (Weininger, 1988). This representation is compact and allows one to use methods developed for fast string comparisons. One can also take advantage of the literature on SMILES-based methods in cheminformatics (Vidal *et al.*, 2005; O’Boyle *et al.*, 2011; Krier and Hutter, 2009). Given this structure representation, we used alignment and scoring methods such as Smith and Waterman (1981) and the Levenshtein distance (Levenshtein, 1966; Damerau, 1964) and looked at the distances between lipids. Building on these distances, one can represent a whole lipidome as a dissimilarity matrix. This numerical representation can then be used for further analyses such as estimating the homology between lipidomes.

Following the use of chemical space models in the field of drug-discovery, the lipid similarity measures were used to define a high dimensional space (Reymond *et al.*, 2010). This approach was evaluated on all lipids of the LIPIDMAPS Structure Database (LMSD) (Sud *et al.*, 2007). Finally, we determined homology between lipidomes of four well characterized yeast strains (Ejsing *et al.*, 2009).

## 2 METHODS

Details of SMILES generation, principal component analysis (PCA), structural similarity methods and annotation of lipids are given in supplementary methods.

### 2.1 Lipid Structure Datasets

Lipid structures for figures 1 and 2 were drawn and SDF files were generated using PubChem Sketcher (Ihlenfeldt *et al.*, 2009). The complete LIPIDMAPS Structure Database (LMSD) in SDF format was downloaded on Nov 9, 2011 from www.lipidmaps.org (LMSDFDownload9Nov11.zip) (Sud *et al.*, 2007).

**Fig. 1.**
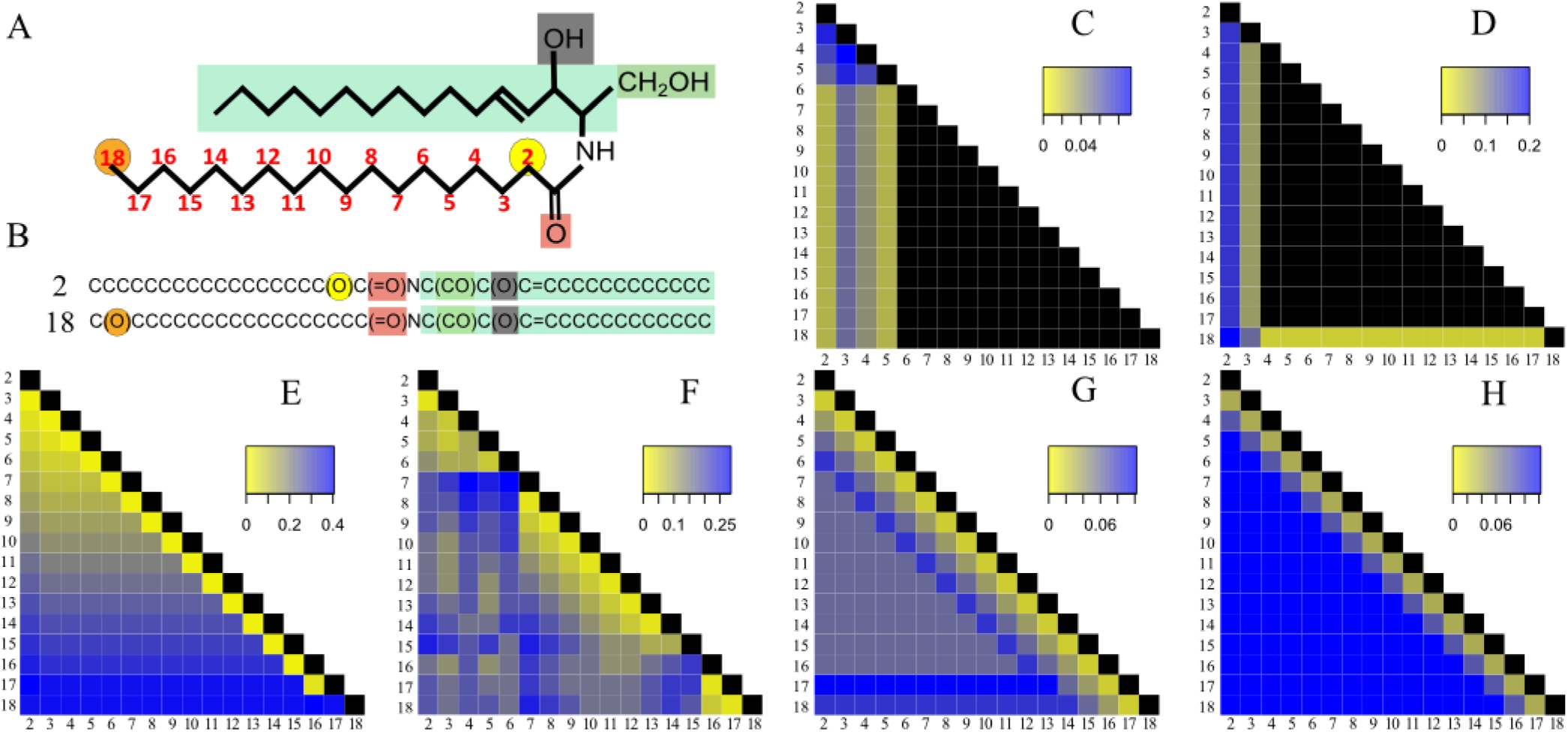
Alignment-based distance calculation algorithms can distinguish isomeric lipid molecules. A. Structure of 17 ceramide molecules consisting of a C16 sphingoid base (light green) and an amide-linked hydroxy fatty acid. The carbon atom number of the hydroxyl group position at the fatty acid chain (red) is used for naming individual molecules. B. SMILES representation of first and last molecules. Color coding of atoms is identical in SMILES-and structure-representations. C-H. Heat map of pairwise distances calculated using Open Babel’s FP2 Fingerprint (C), LINGO (D), Bioisosteric (E), SMILIGN (F), Smith-Waterman (G) and Levenshtein distance (H) algorithms. Bioisosteric method uses CACTVS canonical SMILES whereas for all other methods template-based SMILES were used. Color bars in each panel indicate range of distances values of the particular method. Black denotes a distance value of ZERO, indicating molecular identity. Numbers in rows and columns simultaneously represent the molecule name and the position of hydroxyl group in fatty acid moiety.

**Fig. 2.**
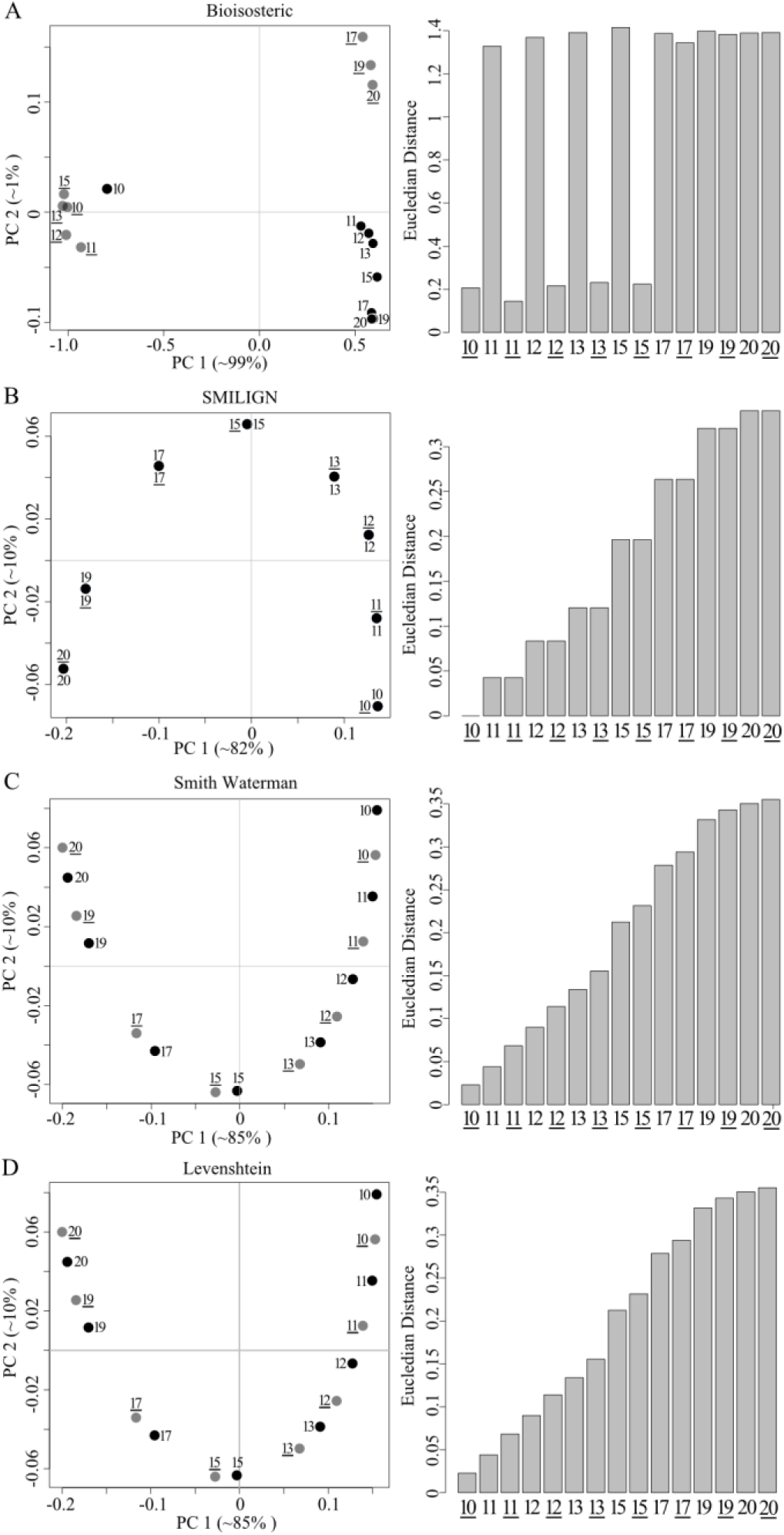
The relationship between phosphatidylinositol (PI) molecules is retained in a two dimensional structural space (The structures and SMILES are provided in Supplementary Result 1). Pairwise distances between 16 PIs were calculated with Bioisosteric, SMILIGN, Smith-Waterman and Levenshtein methods (A-D). PCA was carried out on the distance matrices to generate two-and three-dimensional maps. Approximate contribution of each principal component to the total variance is shown in brackets. Molecules with double bonds are in grey and without double bonds are in black. Euclidian distance between the first molecule PI (10:0/10:0) and all others in the PC1-PC2 plane are shown as bar graphs on the right panels. Molecules were numbered according to the length of the sn2 acyl chain length, wherein an underlined number <u>XX</u> indicate the presence of the double bond.

Lipidome data of yeast mutants was taken from Ejsing *et.al.* (2009). LIPIDMAPS structure drawing tools were customized to draw all required lipid structures for yeast. For ergosterol and ergosta-5,7-dien-3β-ol, SDF files were obtained from the LMSD. SMILES for phytosphingosine 1-phosphate was made by hand from the corresponding phytosphingosine. For some molecules, the number of hydroxylations and double bonds was known, but their position was not. In these cases, a list of possible isomers was generated. The position of double bonds and hydroxylations in yeast fatty acids were taken from previous studies (Hashimoto *et al.*, 2008). Pairwise distances between all isomers were calculated using the Levenshtein distance method (Levenshtein, 1966; Damerau, 1964). The isomer with smallest average distance to other isomers was chosen as representative molecule (Supplementary Result 1).

### 2.2 Template-based SMILES

LIPIDMAPS perl scripts were modified to generate a wider spectrum of lipid structures (Fahy *et al.*, 2007; Sud *et al.*, 2012). These scripts are available at www.lux.fz-borstel.de. Molecule structures were obtained in SDF format and subsequently converted to SMILES using the OpenBabel molecular conversion tool with the default algorithm (O’Boyle *et al.*, 2011). Characters indicating chirality, cis–trans isomerism and charges were removed automatically for the yeast lipidome analysis.

### 2.3 Structural similarity scoring methods

Six similarity scoring methods were tested 1) OpenBabel FP2 Fingerprint 2) LINGO 3) Bioisosteric similarity 4) SMILIGN 5) Smith Waterman Local Alignment 6) Levenshtein distance. Details are given in supplementary methods. The Levenshtein method was applied for analyzing LMSD and yeast lipidome (Figures 4, 5 and 6). This algorithm was originally designed for correcting spelling errors, but the principle can be applied to compare any pair of strings including SMILES (Levenshtein, 1966; Damerau, 1964). The source code used in this study is provided in our website www.lux.fz-borstel.de.

### 2.4 Lipidome Juxtaposition Score (LUX) Calculation

The LUX score is based on the Hausdorff distance (Jain *et al.*, 1999) and summarizes the similarity between lipidomes. In pseudocode, the distance from lipidome A to B is calculated from: for each lipid in A

~~~
find distance d to most similar lipid in B
~~~

~~~
*d*_*sum*_:=*d*_*sum*_+*d*
~~~

~~~
*n* = *n* + 1
~~~

~~~
return *d*_*sum*_/*n*
~~~

This yields the average shortest distance *d*_AB_ from A to B. The larger of *d*_AB_ and *d*_BA_ was used as the lipidome homology score (AB). The LUX score is a simple measure of the homology between a pair of lipidomes. Lower LUX scores signify higher homology.

### 2.5 Hierarchical Cluster Analysis

Complete linkage clustering was performed with R, version 2.14.1, library – ‘stats’ and function ‘hclust’ using the LUX score, Pearson and common lipid count as similarity metrics. An error model for the yeast lipidome analysis was computed by iterating all lipid quantities x of the original data set according to:

~~~
for each lipid with abundance x
~~~

~~~
x′ = x + rnorm(1,0,s)
~~~

~~~
if >x′ > tdetect
~~~

~~~
return x′
~~~

The detection limit tdetect and standard deviation s were defined so that only low abundant lipids were significantly affected. We chose the following three parameter sets: 1) tdetect = 0.003 mol % - 4.3 % of all reported quantities, s = 0.001mol % - 11.4 % of all reported standard deviations 2) tdetect = 0.003 mol %, s = 0.002 mol% - 20.3 % of all reported standard deviations and 3) tdetect = 0.006 mol% - 8.7 % of all reported quantities, s = 0.004 mol% - 34.7 % of all reported standard deviations. The number of occurrences for each branch was counted after 100 iterations using the R library, ape::boot.phylo::prop.part

## 3 RESULTS

### 3.1 Alignment-based similarity scoring methods distinguishes between positional isomers

As a basis for our similarity scoring, we established a template-based SMILES generation algorithm for lipids. We were able to write SMILES in a consistent and predictable manner using template-based structure drawing tools (Fahy *et al.*, 2007; Sud *et al.*, 2012) and OpenBabel default SMILES algorithm (Supplementary Result 2). We then tested alignment methods and distance metrics, analogous to those used for protein or nucleotide sequences. Our quality criterion was based on the methods’ sensitivity to small structural differences commonly found in lipids. The first test dataset consisted of a set of 17 ceramide molecules with the chemical composition C_34_H_68_O_4_N_1_. The position of the hydroxyl group was successively changed from position 2 to 18 at the fatty acid moiety resulting in 17 isomeric molecules (Figure 1A). All isomers were converted into SMILES in which the shift of the hydroxyl group can easily be recognized and we tested six similarity scoring methods (Supplementary Result 2, Figure 1C-H). Three from the literature were used as described under Methods: FP2 Fingerprint (O’Boyle *et al.*, 2011), LINGO (Vidal *et al.*, 2005) and Bioisosteric similarity (Krier and Hutter, 2009). Three methods were introduced here: the SMILIGN, Smith and Waterman (1981) and Levenshtein (1966) distance (Damerau, 1964).

The first clear result is that a large subset of isomeric structures cannot be distinguished by either OpenBabel FP2 Fingerprint or LINGO (Figure 1 C-D). The FP2 Fingerprint algorithm computed a distance of zero for 78 pairs of ceramide isomers (Figure 1C – black pixels). LINGO gave a zero distance for 91 pairs of isomers (Figure 1D). This would only be correct if the molecules were identical. Both methods segment SMILES into shorter sub-strings (1-7 character length in Path-length Fingerprint and 4 characters by LINGO) and apply the Tanimoto coefficient for determining distances. This segmentation into short sub-strings loses the information on the position of the hydroxyl group. In contrast, the Bioisosteric algorithm differentiated all 17 isomeric structures, even though it uses CACTVS Canonical SMILES. There are no zero distances off the diagonal (Figure 1E). The Bioisosteric method also segments SMILES into sub-strings, but in a hierarchical manner, preserving information on the position of the hydroxyl group (Krier and Hutter, 2009). There is a distinct pattern in the heat map of the Bioisosteric method characterized by a gradual increase in distance values for isomers having the hydroxyl group closer to the terminal methyl carbon. The Bioisosteric method returns a distance of 0.13 units for the shift of the hydroxyl group from position 5 to 7 (Figure 1E - light green pixel), but returns 0.26 units for position 10 to 12 (Figure 1E - yellow pixel) and for positions 16 to 18, a distance of 0.41 was calculated (Figure 2E - white pixel). This dependence of the distance values on the position of the hydroxyl group leads to an unwanted weighting which is a clear problem with the approach.

In the SMILIGN algorithm, SMILES strings are treated as if they were amino acid sequences and a multiple sequence alignment was calculated with MUSCLE (Edgar, 2004). The pairs of lipids were rescored using an identity matrix. The SMILIGN method distinguished all 17 ceramide isomers (Figure 1F), but we noticed an irregular distribution of distance values. For example, comparing molecule pairs where the hydroxyl group was shifted by one position 11-12, 12-13, 13-14 and 14-15 resulted in four different distance values of 0.03, 0.13, 0.25 and 0.06 units respectively. In this regard, we identified two problems with the algorithm. First, there were several misalignments that lead to incorrect distances. Second, one needs 35 characters to represent all the structural details of all lipid molecules of the LMSD (LipidMaps Structure Data-base). The software is limited to only 20 characters and too much information is lost. To overcome these two limitations of SMILIGN, we tested two pair-wise alignment methods that do not require conversion to amino acid sequences.

With the Smith-Waterman method, pair-wise alignments are carried out directly with the SMILES strings. All ceramide isomers were distinguished, but we noticed an anomaly in distance values for molecules 17 and 18 (Figure 1G). A closer examination of the pair-wise alignments revealed an inherent issue when applying local alignment procedure to lipids. In the aligned SMILES pairs 2-17 and 2-18, the hydroxyl groups in the fatty acid were ignored, while for the pairs 2-15 and 2-16 the characters were retained. The Smith-Waterman algorithm is designed to find high scoring regions in strings, so differing ends are ignored by design and not by accident. This means that functional groups at the omega position are ignored, despite their role in biology (Kniazeva *et al.*, 2004). Although one could try to adjust parameters, the Smith-Waterman method is fundamentally not appropriate for this kind of comparison.

Finally, we tested the Levenshtein distance for measuring similarities between lipid molecules (Figure 1H). Unlike Smith-Waterman, the Levenshtein approach always aligns all characters for a given pair of SMILES. This method was the most successful. It distinguished all ceramide molecules and for each molecule, it correctly scored and ranked distances up to the molecule’s third closest isomers. From the fourth closest isomer onwards, a fixed distance of 0.12 was determined. Unlike other methods, it guarantees a symmetric distance matrix with no unwanted weighting of groups due to their positions.

These tests of structural similarity measures led to two conclusions. First, the alignment step is necessary. Second, the Levenshtein distance was most likely to be generally applicable for all molecules in a lipidome.

### 3.2 From structural similarity to chemical space

A set of distances between *n* molecules defines an (*n* −1) dimensional space. The coordinates of molecule *i* are simply the distances to all members of the set (including the zero distance to molecule *i* itself). This is formally a vector space so similar molecules will have similar coordinates. It is, however, not very compact and because of structural similarities, coordinates in some dimensions would be highly correlated with others. Principal component analysis (PCA) was then used to reduce the dimensionality and see how much information would be lost. The first test was performed on a set of 16 phosphatidylinositol molecules (Supplementary Result 1).

Considering just the first two principal components was sufficient to highlight problems with some of the distance measures. For example, the map in Figure 2A shows a clear weakness with the Bioisosteric method. The extension of the fatty acid chain at the sn2 position and degree of unsaturation are not accurately rep resented (Figure 2A, scatter plot). We also computed the Euclidian distance between molecules in the plane of the first two components. This showed an inconsistent trend in the distance increase with each structural alteration (Figure 2A, bar graph). Principal components can often be interpreted in terms of the original descriptors and in the case of SMILIGN, the first two components are dominated by the extension of the acyl chain at the sn2 position (Figure 2B). For SMILIGN, the first two principal components are not sufficient to distinguish molecules that differ only in the presence of a double bond, but the third principal component does capture it (Supplementary Result 3).

In contrast, distances based on Smith-Waterman and Levenshtein algorithms reflected all gradual structural changes in the molecules (Figure 2 C,D). In both cases, the projection leads to a set of points in a ‘U’ shape and, if we take molecule 10 as a reference, stepwise changes to the chemistry are reflected in distinct shifts in the principal coordinates. We further recognized that the changes in coordinates, when the acyl chain is extended by two methylene groups (molecules 15-17, 17-19) are about twice as large as the difference between pairs differing by a single methylene group (Figure 2C-D). The first two principal components combined, accounted for 95% of the variability in the underlying data set for Smith-Waterman and Levenshtein. Summarizing the results, we see the Levenshtein method coupled with template-based SMILES as the best approach for calculating structural differences in small molecule sets. PCA is an appropriate way to reduce dimensionality and the relation between molecules can be depicted in a PCA map, which we treat as chemical space.

The set of 16 phosphatidylinositol molecules is useful for highlighting details, but one is interested in using the method on much larger molecule sets. To this end, we used the 3510 sphingolipids (SP) from LMSD as a test dataset (Sud *et al.*, 2007). All lipid structures were converted into template-based SMILES and pairwise distances were computed using the Levenshtein method. Figure 3A shows the position of all molecules in terms of the first three principal components. There are two clear observations. First, three principal components account for 99% of the total variance and no two SP have the same coordinates. Second, there was no biochemical knowledge put into the procedure, but the molecules cluster naturally into chemically similar groups (Figure 3A). Sphingosines, ceramides and phosphosphingolipids were clustered separately from the complex glycosphingolipids (GSL). Furthermore, the acidic and neutral GSL where placed in different clusters. Looking at the globo, lacto, neolacto and isoglobo series of neutral GSL, one can see changes in the sugar moiety and a clear separation from the simple ‘Glc’ series (Figure 3B). This observation fits the intuitive expectation that the ‘Glc’ series with simple sugar moieties (glucose, galactose or lactose) should be farther from lipids with complex sugar composition. We noted that changes in the sugar moiety composition of neutral GSL, which have a strong impact on biochemical behavior, were separated by a larger distance compared to changes in the ceramide backbone (Figure 3B). In addition, we were intrigued by the recurring appearance of geometrical patterns in the form of ‘I’, ‘C’ and ‘L’ shapes and investigated the structure within these clusters. Within each cluster, lipids were organized based on changes in the ceramide moiety (Figure 3C) so that, for example, eight molecules of the isoglobo series formed a twisted ‘L’ shape with each successive lipid carrying a gradual change in the ceramide backbone (Figure 3C – light brown colored points). Analogous geometric arrangements were observed for the globo, lacto and neolacto series (Figure 3C – red, violet and light-blue points). Next we tested, how all the 30 150 lipids of the LMSD would be organized in a chemical space based on only the structural similarity. All lipid molecules had unique coordinates in the computed chemical space, indicating that our approach can distinguish between all lipid structures within known, natural lipidomes. With no additional input, the method grouped lipids into clusters that correspond to the popular lipid classification of LIPIDMAPS (Figure 4A) (Fahy *et al.*, 2008).

**Fig. 3.**
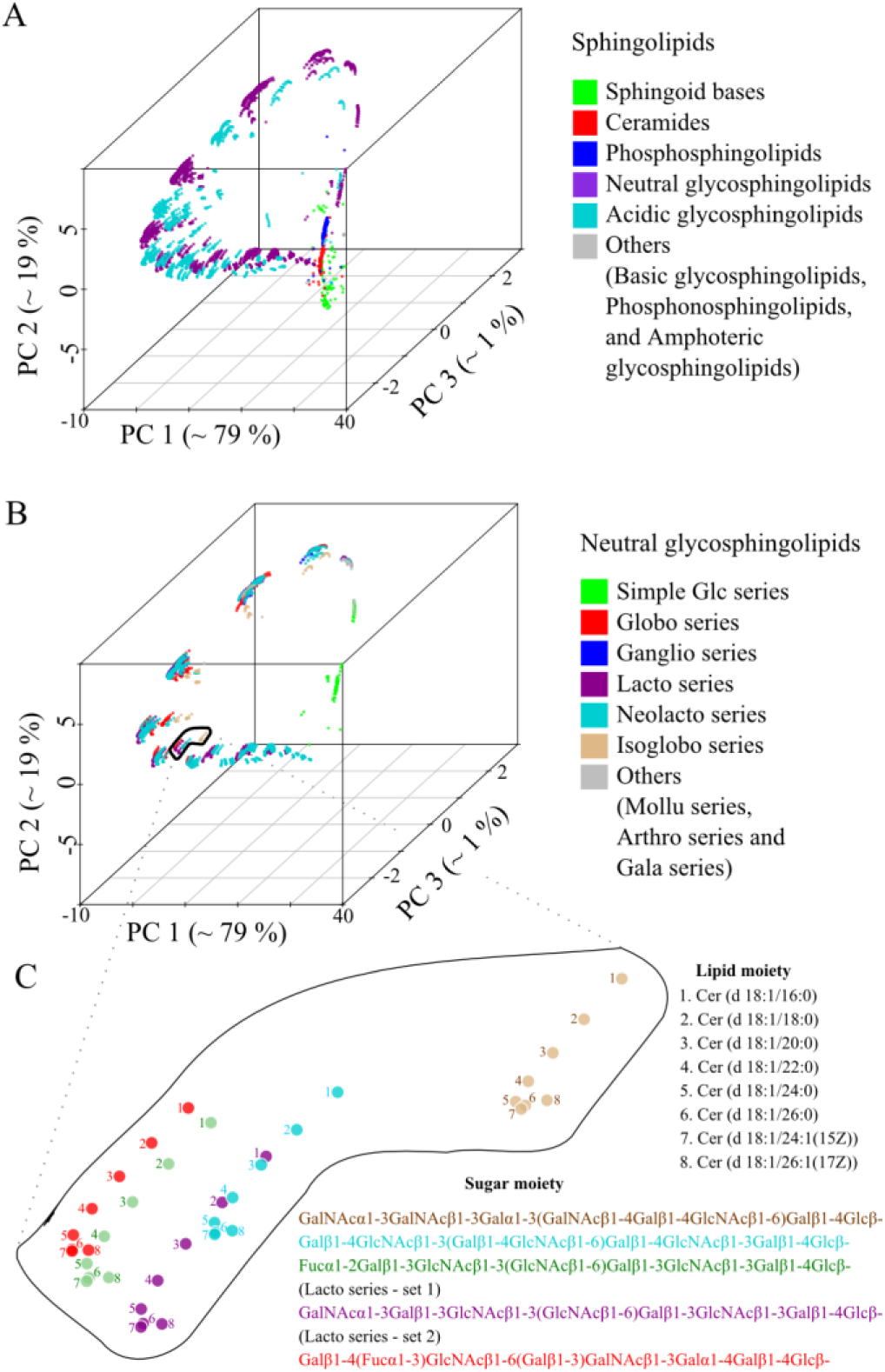
The structural space model clusters thousands of sphingolipids (SP) according to their chemical relationship. A. Three dimensional map of 3510 SP obtained by PCA from a pair wise distance matrix calculated with Levenshtein distance. B. Plot of all neutral SP within the same coordinate system as panel A indicating several associated glycosphingolipid series. C. Complex glycosphingolipids are highlighted showing the influence of structural changes in the ceramide backbone and sugar moiety.

**Fig. 4.**
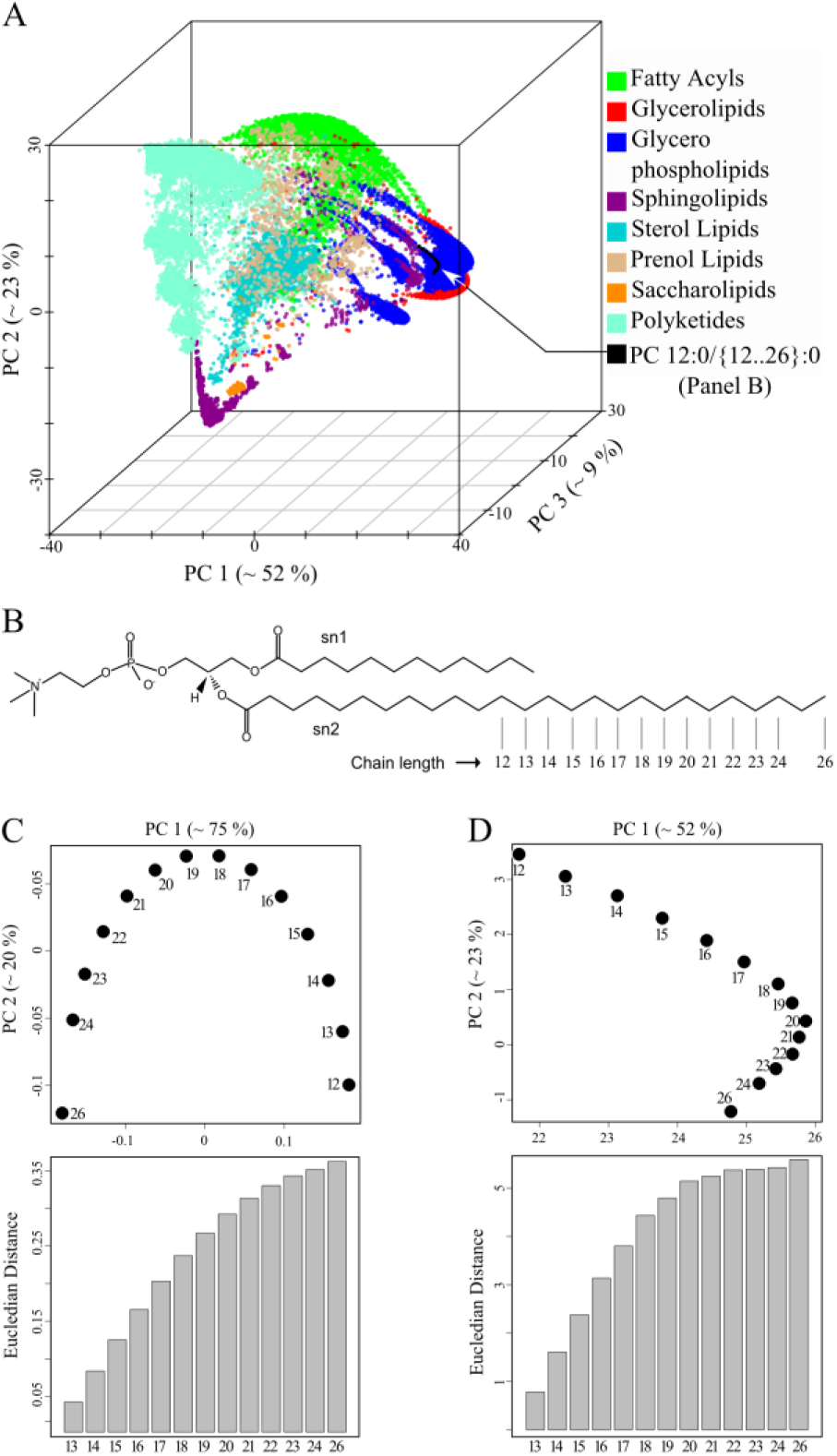
Spatial distribution of related phosphatidylcholines (PC) moelcules remains stable in the background of large structure data sets. A. Lipid map of 30 150 lipid molecules obtained from LMSD. Pair wise distances were calculated using the Levenshtein method of template-base SMILES. B. Structures of the 14 PC molecules. Molecules are named based on the number of carbon atoms of the sn2 acyl chain. C. Two dimensional map of the selected PC molecules displaying their chemical relation to each other. Euclidian distances in PC1-PC2 plane between the smallest PC (12) and all others are shown in the bar graph. D. Spatial distribution of 14 PCs in the background of 30136 lipids determined from three principal components and projected on the PC1-PC2 plane. Euclidian distances between first molecule (12) and all other was determined form the first two principal components and its trend is shown as bar graph.

Fatty acyls, glycerophospholipids (GPL), sphingolipids (SP) and polyketides occupied opposite ends of the chemical space. In contrast, glycerolipids and GPL shared a common region because of their head group similarity. Sterol lipids formed a distinct cluster due to their unique four-ringed core structure. Prenol lipids were widely distributed in the chemical space reflecting their varying chemical composition. For GPL, we observed several distinct clusters, which on closer examination, could be recognized as spatially separated lipid classes like phosphatidylcholine (PC), phosphatidylserine (PS) and phosphatidylinositols (PI).

### 3.3 The spatial organization of lipids is robust to changes in background molecule ensemble

As with the set of PI molecules described above, (Figure 2D) the PC molecules in the two-dimensional representation form an inverted ‘U’ pattern (Figure 4C). However, the PC molecules formed a flipped ‘L’ pattern if all other 30 136 lipids of the LMSD were present (Figure 4D). In both cases, the sequential arrangement of the PC-molecules in the two-dimensional chemical space accurately represents the gradual increase in acyl chain length. We also observed a gradual increase of the Euclidian distance from the molecule PC (12:0 / 12:0) to all species with longer sn2 acyl chains (Figure 4 C, D). When we gradually increased the complexity of the set of lipid molecules, we noticed that the PCA approach could disturb relationships between structurally similar molecules. In the case of a set consisting of only GPLs and only GPLs and SPs (data not shown), we noticed that the distances between molecule 12 and molecules 21-26 did not reflect the sn2 chain length increase anymore. Interestingly, one can observe that the gradual addition of more diverse lipid structures spanning a broader chemical variety compensates for this bias. It seems that the Levenshtein distances and the projection to a chemical space automatically reconstructs conventional lipid class definitions. The next natural step is to use these lipid coordinates to analyze and compare complete lipidomes.

### 3.4 The Lipidome jUXtaposition (LUX) score, a single metric for calculating the homology between lipidomes

The approach to lipidome comparison was then tested on real data. All lipids from four yeast strains BY474, Elo1, Elo2 and Elo3 (Ejsing *et al.*, 2009) were combined, yielding a reference lipidome with 248 members, each with unique coordinates (Supplementary Result 1). For clarity, this is shown in a 2D map (Figure 5A), which is the basis of comparisons of the four strains and two culturing temperatures (24°C and 37°C). Triacylglycerols (TAG) occupy the largest area on the map in terms of the variety of structures. Mannose-bis(inositolphospho)ceramides (M(IP)_2_C) form a distinct cluster located in the top-left quadrant of the reference map. In the top right quadrant of the reference map, there is a cluster of GPLs consisting of phosphatidic acid (PA), phosphatidylethanolamines (PE), phosphatidylcholines (PC) and TAG. The reference lipidome map clearly shows temperature-and strain-specific changes. The lipidomes of the wild type yeast strains BY4741 and Elo1 grown at 24°C showed only minor differences (Figure 5B). In contrast, the lipidome of the Elo2 mutant is very different to the wild type strain BY4741 (Figure 5C). The mutation has led to dramatic changes amongst the inositol phosphorylceramides seen in the top-left quadrant and the appearance of new species not present in the wild type. Using this well-defined lipidome map, one can determine the closest related lipid in the wild type strain If one calculates the distances that lipids would have to move to make the members of each pair overlap, one can use the Hausdorff distance to compare lipidomes (arrow marked lipids, Fig 5C, D). For that, we chose the coordinates of all lipids in the two dimensional coordinate system of the first lipidome and determined the Euclidean distance to its closest structural neighbor in the second. Subsequently, the average of all distances was determined, including all distance values of zero for identical molecular species. Because the Hausdorff distance depends on the direction of the comparison, we used the maximum of the two values (max(*d*_*AB*_,*d*_*BA*_)). We named this measure as the ‘Lipidome jUXtaposition (LUX) score’. This score is a distance, so larger values indicate more dissimilarity and identity results in a LUX score of zero. From that perspective, one can see that the LUX score between BY4741 and the Elo2 strain is three-fold larger than the distance between BY4741 and Elo1 (Figure 5B, C).

**Fig. 5.**
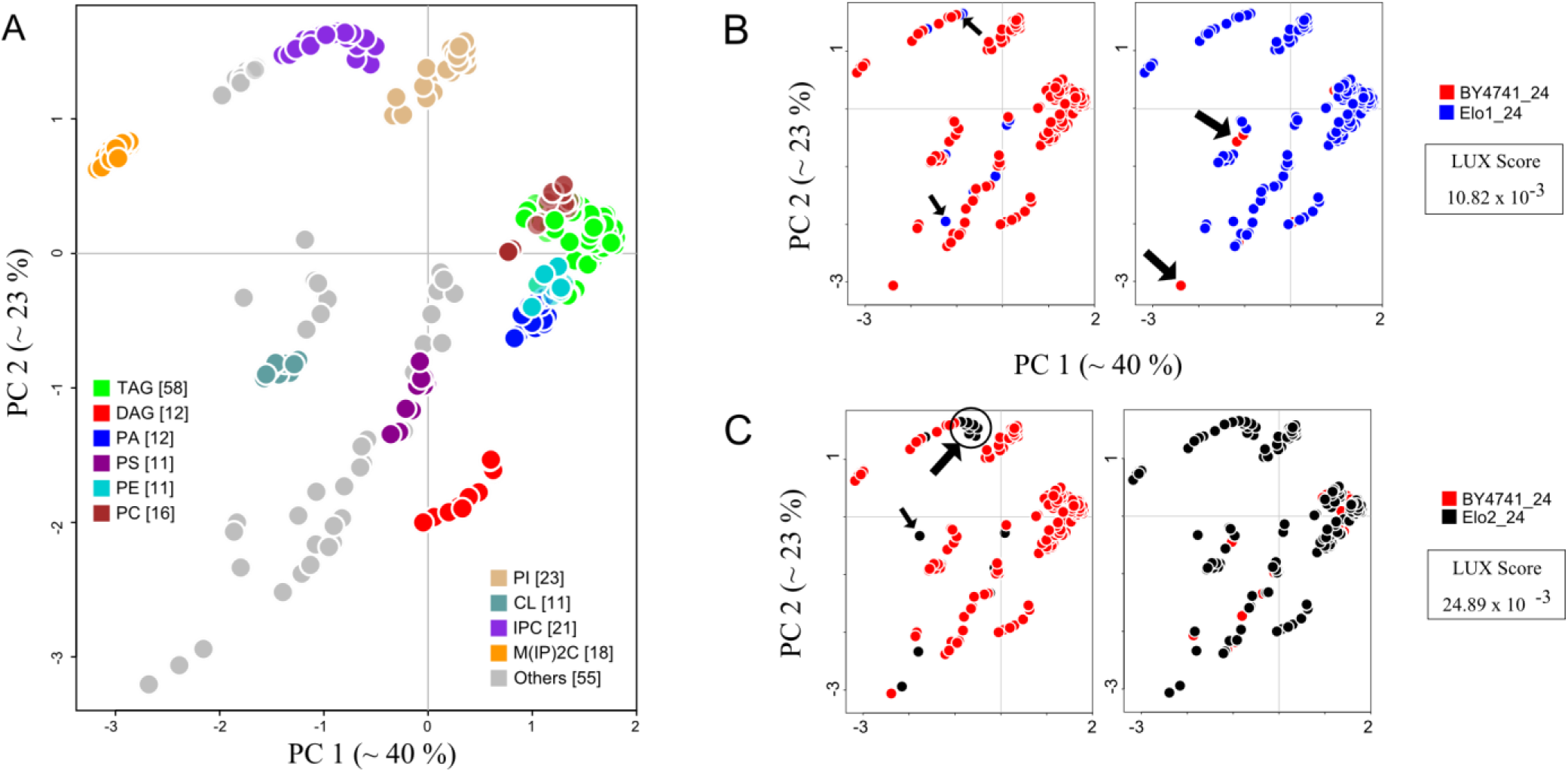
Lipidome maps highlight relationships between yeast strains. A. All lipids from yeast strains, BY4741, Elo1, Elo2 and Elo3 cultured at 24°C and 37°C are combined to create a reference map of the yeast lipidome. Each colored circle corresponds to a unique lipid. B. Comparison of lipidomes from strains BY4741 and Elo1 (cultured at 24°C). Arrows in first plot indicate lipids that are present in Elo1, but not in BY4741 and vice versa in the second plot. C. Comparison of BY4741 and Elo2 lipidomes. A two dimensional Lipidome jUXtaposition (LUX) score is calculated for a pair of lipidomes using reference-map coordinates (Supplementary Result 1).

Next we evaluated the LUX score by computing a hierarchical clustered tree of all eight reported lipidomes of yeast (Figure 6A) and compared it to dendrograms based on the lipid concentrations (Figure 6B), and simply by counting common lipids (Figure 6C). That allowed us to test if our approach can correctly depict the genetic and phenotypic relationship between the yeast strains reported earlier (Ejsing *et al.*, 2009). The tree computed from the LUX score as well as common lipid count indicates that mutation of the Elo1 gene had less influence on the composition of the lipidome than the temperature shift. Both strains, BY4741 and Elo1 were closest neighbours to each other at the culturing temperature of 24°C and 37°C. The lipidomes from mutant strains Elo2 and Elo3 were clustered together using the LUX score (Fig. 6A) but in counting common lipids, Elo2 clustered with BY4741 and Elo1 (Fig 6C). This marks the major difference between both metrics. It was reported that no aberrant phenotype for Elo1 was observed and that Elo2 and Elo3 had distinct alterations in their intracellular organization (Ejsing *et al.*, 2009; Oh *et al.*, 1997), which seems better represented with the LUX score. However, we verified this finding with an error model that modify only the presence and quantity for low abundant lipids to estimate a robustness for the observed clustering. One can recognize that the LUX score (Fig 6A) as well as the common lipid count (Fig 6C) comprise a sufficiently robust tree topology and groups Elo2 systematically different. We concluded from this experiment that compositional differences itself are useful to assign a phenotype while comparison purely based on quantities are dominated by changes of abundant lipids (Fig 6B). We also note that just counting of lipids is a simplistic, binary measure of compositional differences. In contrast, the LUX score provides a refined measure of lipidome structural diversity, which we recognize as an advantage.

**Fig. 6.**
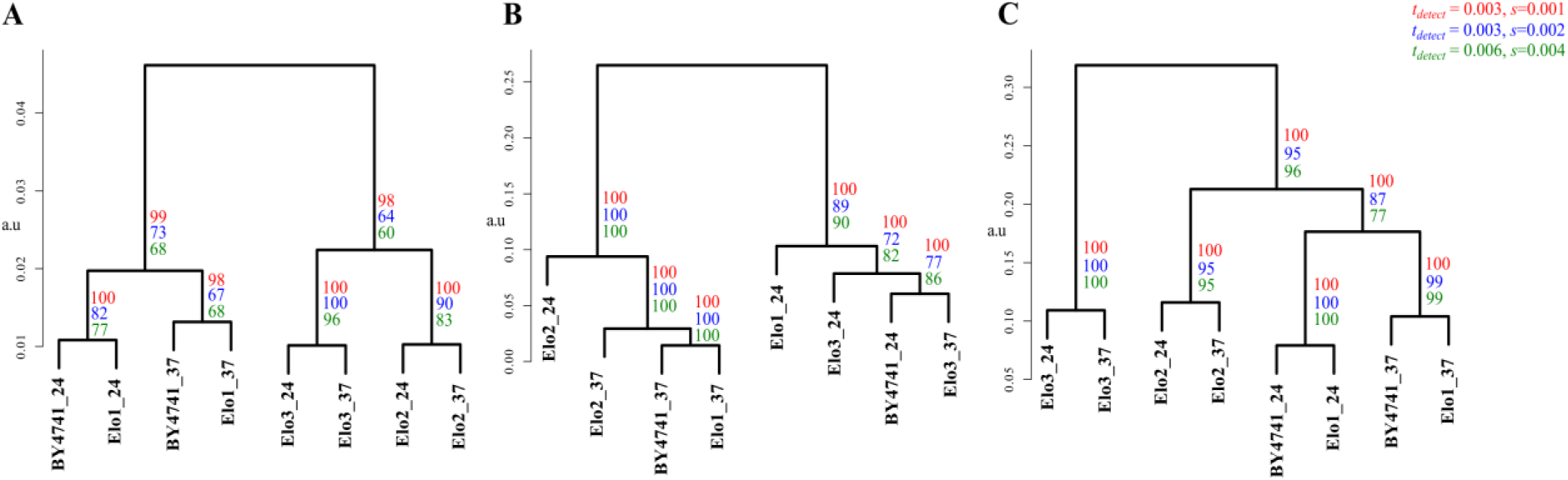
For the yeast strains, BY4741 (wild type), Elo1, Elo2 and Elo3 (*Elongase* mutants) cultured at 24°C and 37°C, dendrograms were computed from two-dimensional LUX scores (A) Comparing concentrations of common lipids (B) and counting the percentage of common lipids in a pair of lipidomes (C). All dendograms are based on complete linkage using Euclidean distance as the similarity metric (a.u - arbitrary units). The number of occurrences for each branch in 100 iterations is indicated with coloured numerals that correspond to the utilized parameter set for detection threshold *t*_*detect*_ and standard deviation *s* applied in the error model (see methods).

The complexity of the yeast lipidome comparison is relatively small compared to higher organisms. Nevertheless, the two-dimensional structural space reflecting 63% of the overall variability of the dataset (Fig. 5A) is sufficient to determine lipidome similarities based upon the LUX score. We also note that the tree topology does not change whether one uses just three principal components (covering 83% of the variability) or the original pairwise distance matrix (data not shown). This indicates that our approach enables a simple way to reduce the complexity of large lipid structure datasets, which can further help to depict results of a lipidome homology analysis in a well-defined manner. In this way researchers can report their findings for interspecies, cell type and cell compartment lipidome analyses in a defined graphical representation. At the same time, the LUX score determination workflow can be customized with regards to the complexity of the lipidome study.

## 4 DISCUSSION

Our study offers a general approach to characterizing and comparing lipidomes based on the structures of their constituents. It is certainly useful for making function / phenotype associations and allows one to correlate changes with habitat, genetic relationships and environmental stresses. The approach is dependent on the initial SMILES strings which is both an advantage and possible weakness. One can consider a comparison with small molecule classification. There, the problem is sometimes easier, especially when one is dealing with derivatives which are closely related, but even in small molecule chemoinformatics, there is no universally accepted scheme (Bender et al, 2009). Optimization of such structures often depends on the interaction sites of a protein and pharmaceutical requirements for administration of drugs (Mohanapriya and Achuthan, 2012; Ahmed and Ramakrishnan, 2012). In this study, the analysis does not stop after comparing the details of individual structures. The larger aim is whole lipidome comparison and these are sets of structures whose members are functionally related. In this work, we leverage a SMILES generation scheme which works well on large sets, but there will probably be pathological examples where it does not perform well. It definitely seems useful when working with lipids where it reflects 1) chain length 2) double bond position and 3) bond frequency. However, lipids are very special with regards to their structural diversity, and some better similarity metrics might be available in future.

The definition of structural similarity and chemical space model also concisely depicts the complexity of a lipidome. The projection down to two-and three-dimensional maps lead to clusters which fit standard lipid nomenclature. This means that one can intuitively see qualitative differences between lipidomes. The reference map for multiple comparisons also shows changes in the overall organization of a lipidome which can support functional association re lated to membrane organization and metabolic adaptations. For the analysis of compositional differences between lipidomes and its interpretation, we recommend to apply an error model as introduced in this study. We recognized that clustering approaches are often not verified with an error model, which negatively affects the value of subsequently derived biological and/or medical interpretations. The lipidome comparisons in this study are solely based upon an identity matrix for exchange values which does not account for quantitative changes. This is parsimonious, but obviously not optimal for comparing biological systems in terms of their lipidomes. In future work, we will test how quantitative changes should be weighted with respect to structural changes. We will estimate such weight measures from well understood model systems based on larger data sets that are now becoming available (Voynova *et al.*, 2014; Tarasov *et al.*, 2014). However, this study shows that the structural composition of a lipidome is sufficient to recognize the degree of genetic alteration and growth temperature dependence in yeast strains in an unsupervised manner. In contrast, all correlation based methods using lipid quantities as input failed.

The growth in experimental data combined with methods like the LUX score may provide a basis for disease and trait association studies as used in genome research.

## ACKNOWLEDGEMENTS

The authors are thankful to Eli Lebow, Geoff Hyde, Mukund Thattai, Sandeep Krishna and Kurt Fellenberg for critical discussions.

*Funding*: This work was supported by a Wellcome Trust India Alliance Senior Fellowship to DS and by funds of the German Research Foundation of the SFB-TR22 consortia. CM was supported by UGC Junior Research Fellowship and NCBS-TIFR Graduate Student Fellowship.

